# A photostable monomeric superfolder GFP

**DOI:** 10.1101/811588

**Authors:** Fernando Valbuena, Ivy Fizgerald, Rita L. Strack, Neal Andruska, Luke Smith, Benjamin S. Glick

**Affiliations:** Department of Molecular Genetics and Cell Biology The University of Chicago 920 East 58th St. Chicago, IL 60637

## Abstract

The green fluorescent protein GFP from *Aequorea victoria* has been engineered extensively in the past to generate variants suitable for protein tagging. Early efforts produced the enhanced variant EGFP and its monomeric derivative mEGFP, which have useful photophysical properties, as well as superfolder GFP, which folds efficiently under adverse conditions. We previously generated msGFP, a monomeric superfolder derivative of EGFP. Unfortunately, compared to EGFP, msGFP and other superfolder GFP variants show faster photobleaching. We now describe msGFP2, which retains monomeric superfolder properties while being as photostable as EGFP. msGFP2 contains modified N- and C-terminal peptides that are expected to reduce nonspecific interactions. Compared to EGFP and mEGFP, msGFP2 is less prone to disturbing the functions of certain partner proteins. For general-purpose protein tagging, msGFP2 may be the best available derivative of *A. victoria* GFP.

## 1 INTRODUCTION

The green fluorescent protein (GFP) from *Aequorea victoria* was the first fluorescent protein (FP) to be used as a reporter and as a tool for labeling proteins, cells, and tissues (1). Multiple rounds of protein engineering have yielded improved GFP variants. Initial efforts focused on photophysical properties, and led to creation of the enhanced variant EGFP (1). Compared to wild-type GFP, EGFP shows faster chromophore maturation as well as a red-shifted excitation spectrum that allows fluorescence to be excited with blue light. EGFP is bright and photostable, and has been the most widely used GFP variant.

Yet EGFP and other first-generation GFP variants have limitations. Most notably, GFP has a weak tendency to dimerize, and this property can induce artifactual clustering of tagged proteins in the crowded environment of a cell (2, 3). The self-association of GFP variants is effectively eliminated by point mutations that disrupt the dimerization interface (4). For example, an A206K mutant of EGFP yields the monomeric variant mEGFP. A more subtle issue is that GFP can misfold to form nonfluorescent aggregates. For example, when EGFP is fused to bacterial proteins that form inclusion bodies, the EGFP tag fails to become fluorescent. This observation was the basis for creating a “superfolder” GFP that achieves the fluorescent state even in inclusion bodies (5). Superfolder GFP can outperform other GFP variants, most notably in oxidizing environments such as the lumen of the endoplasmic reticulum or the periplasm of gram-negative bacteria (6, 7). The original superfolder GFP contained an A206V mutation (5), but superfolder behavior is also obtained with the monomerizing A206K mutation (3), thereby capturing the monomeric and superfolder properties in a single variant. We previously used a monomeric superfolder GFP called msGFP as a model protein to track secretion in yeast (8).

A disadvantage of currently available superfolder GFP variants is that they photobleach faster than EGFP or mEGFP (5). This concern limits the utility of superfolder GFP variants for live-cell imaging. Our goal was to create a new monomeric superfolder GFP that would retain the desirable photophysical properties of EGFP.

As part of this engineering effort, we also explored an issue that has received little attention, namely, the peptides at the N- and C-termini of FPs. Our interest in this topic was prompted by earlier work to optimize DsRed, a tetrameric red FP. DsRed undergoes higher-order aggregation that can be suppressed by mutating residues near the N-terminus (9-11). Furthermore, mutations near the N-terminus of DsRed enhanced bacterial expression (10). A different approach was taken for generating the monomeric DsRed variant mCherry, which contains N- and C-terminal peptides derived from EGFP (12). The rationale was that EGFP is a well-behaved FP, so its N- and C-terminal peptides were expected to confer similarly good behavior on other FPs. This practice of incorporating the terminal peptides of EGFP has become widespread when developing new FPs. However, we report here that the terminal peptides from EGFP can cause bacterial cytotoxicity when present in FPs such as mCherry. To avoid this problem, we incorporated alternative N- and C-terminal peptides into msGFP2.

The enhanced utility of msGFP2 as a fluorescent tag will be most apparent with a subset of partner proteins. We demonstrate that msGFP2 can outperform EGFP or mEGFP in certain protein fusions.

## 2 RESULTS

### 2.1 Introduction of point mutations into EGFP

We previously modified EGFP using the following approach (8). First, we made the eight superfolder mutations S30R, Y39N, F99T, N105T, Y145F, M153T, V163A, and I171V. Those mutations are identical to the ones in the original superfolder GFP (5), except that F99T was chosen instead of F99S because in the closely related GFP homolog phiYFP (13), threonine is present at the position corresponding to F99. Second, instead of the A206V mutation in the original superfolder GFP, we made the monomerizing mutation A206K (3, 4). Third, to revert the accidental introduction of leucine at position 231 during the creation of EGFP (1), we made an L231H mutation. Collectively, these changes generated a monomeric superfolder variant of GFP.

### 2.2 Replacement of the terminal peptides in mCherry and mScarlet-I

The next set of changes focused on the N- and C-terminal peptides. This effort was motivated by our analysis of mCherry, which contains N- and C-terminal peptides derived from EGFP (12). We found that expression of mCherry in the *E. coli* strain DH10B results in cytotoxicity, presumably because mCherry associates with itself or with other cellular components. Screens of mCherry mutant libraries implicated the termini of the polypeptide chain in cytotoxicity (data not shown). Therefore, to optimize N- and C-terminal peptides for use in an improved GFP variant, we employed mCherry as a test protein.

The six N-terminal amino acids of wild-type GFP are MSKGEE. In EGFP, immediately after the start codon, a valine codon (V1a) was inserted to create a Kozak sequence for efficient mammalian expression (1), so the N-terminal peptides of EGFP and mCherry are MVSKGEE. We replaced the N-terminal peptide of mCherry with MDSTES, which incorporates the DST tripeptide that was found to enhance the solubility and bacterial expression of DsRed derivatives (10). The V1a codon was omitted because the D2 codon creates a Kozak sequence.

The six C-terminal amino acids of wild-type GFP and EGFP are MDELYK. Because this peptide can potentially undergo electrostatic and hydrophobic interactions, we replaced the C-terminal six residues of mCherry with the linker sequence GSSGSS. At the same time, the previous three amino acids were changed from TGG to GSQ to match the sequence in another monomeric DsRed variant termed DsRed-Monomer (14). The resulting variant with modified N- and C-termini was termed mCherry2B (15). Subsequently, to match the more standard practice of using glycine-rich linkers (16), we switched to using GGSGGS for the C-terminal peptide. A variant containing MDSTES at the N-terminus and GGSGGS at the C-terminus was termed mCherry2C.

Figure S1A shows expression in DH10B cells of mCherry, mCherry2C, and mCherry variants containing either the N-terminal MDSTES replacement alone or the C-terminal GGSGGS replacement alone. Compared to the colonies expressing mCherry, the colonies expressing mCherry2C were substantially larger. Similar results were obtained with mCherry2B (data not shown). The N-terminal replacement evidently made the major contribution to reducing cytotoxicity. This effect is probably not due to lower expression because the MDSTES coding sequence is expected to increase bacterial expression (10). We conclude that replacing the terminal peptides of mCherry can reduce cytotoxicity.

To determine whether this improvement extends to another FP, we examined mScarlet-I, a synthetic monomeric red fluorescent protein with N- and C-terminal peptides from EGFP (17). The N-terminal peptide of mScarlet-I was replaced with MDSTEA because we found that MDSTES diminished the fluorescence (data not shown), and the C-terminal peptide was replaced with GGSGGS, yielding a variant that we termed mScarlet2-I. Although the difference between mScarlet-I and mScarlet-2I was less pronounced than the difference between mCherry and mCherry2C, the DH10B colonies expressing mSclarlet-2I were larger on average than those expressing mScarlet-I (Figure S1B,C and Figure 1). The combined results indicate that the terminal peptides of EGFP can render some FPs cytotoxic.

**FIGURE 1.**
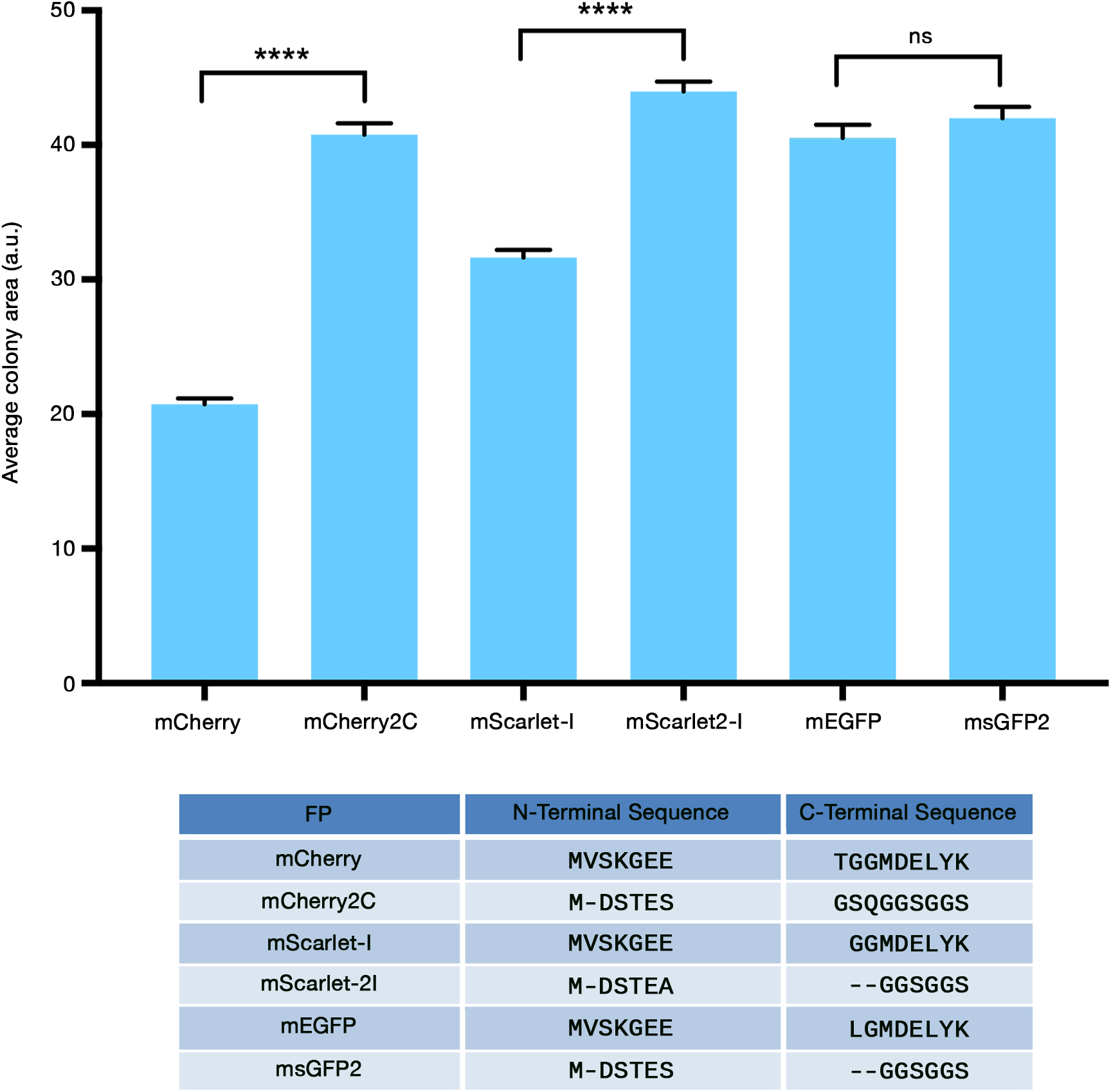
FP cytotoxicity in bacteria can be reduced by modifying the protein termini. *E. coli* cells of strain DH10B were transformed with the indicated FP expression constructs, and colony areas were measured as described in Methods. Average colony areas are plotted in arbitrary units. Bars indicate SEM. Statistical significance was measured using Welch’s *t*-test. Four asterisks indicate significance at P < 0.0001, and “ns” indicates no significant difference. The table at the bottom lists the N- and C-terminal peptides of the FPs, with the dashes indicating amino acids that are missing from one of the two FPs in each pair.

### 2.3 Replacement of the terminal peptides in monomeric superfolder GFP variants

To generate the original msGFP, we modified our monomeric superfolder GFP variant to contain the N-terminal peptide MDSTES and the C-terminal peptide SSGSSG (8). The MDSTES sequence preserves a single glutamate, which is needed at position 5 or 6 of GFP to obtain fluorescence (18). During further engineering to generate msGFP2, the C-terminal eight residues were replaced with the glycine-rich peptide GGSGGS, and an additional photostabilizing mutation was introduced as described below.

When expressed in DH10B cells, mEGFP (which contains the EGFP terminal peptides) yielded colonies as large as those obtained with msGFP2 (Figure S1B,C and Figure 1). Thus, for unknown reasons, cytotoxicity due to the EGFP terminal peptides was apparent with two monomeric red FPs but not with a monomeric GFP. We nevertheless included the modified N- and C-terminal peptides in msGFP2, based on the concern that even for a GFP variant, the EGFP terminal peptides might have deleterious effects under some circumstances.

### 2.4 Generation of the photostable msGFP2 variant

Superfolder GFP variants have been reported to photobleach more quickly than EGFP (5, 19), and we observed this effect as well. When purified msGFP and EGFP were illuminated with blue light on a microscope stage (Figure 2A), msGFP (light green curve) was substantially less photostable than EGFP (dark blue curve).

**FIGURE 2.**
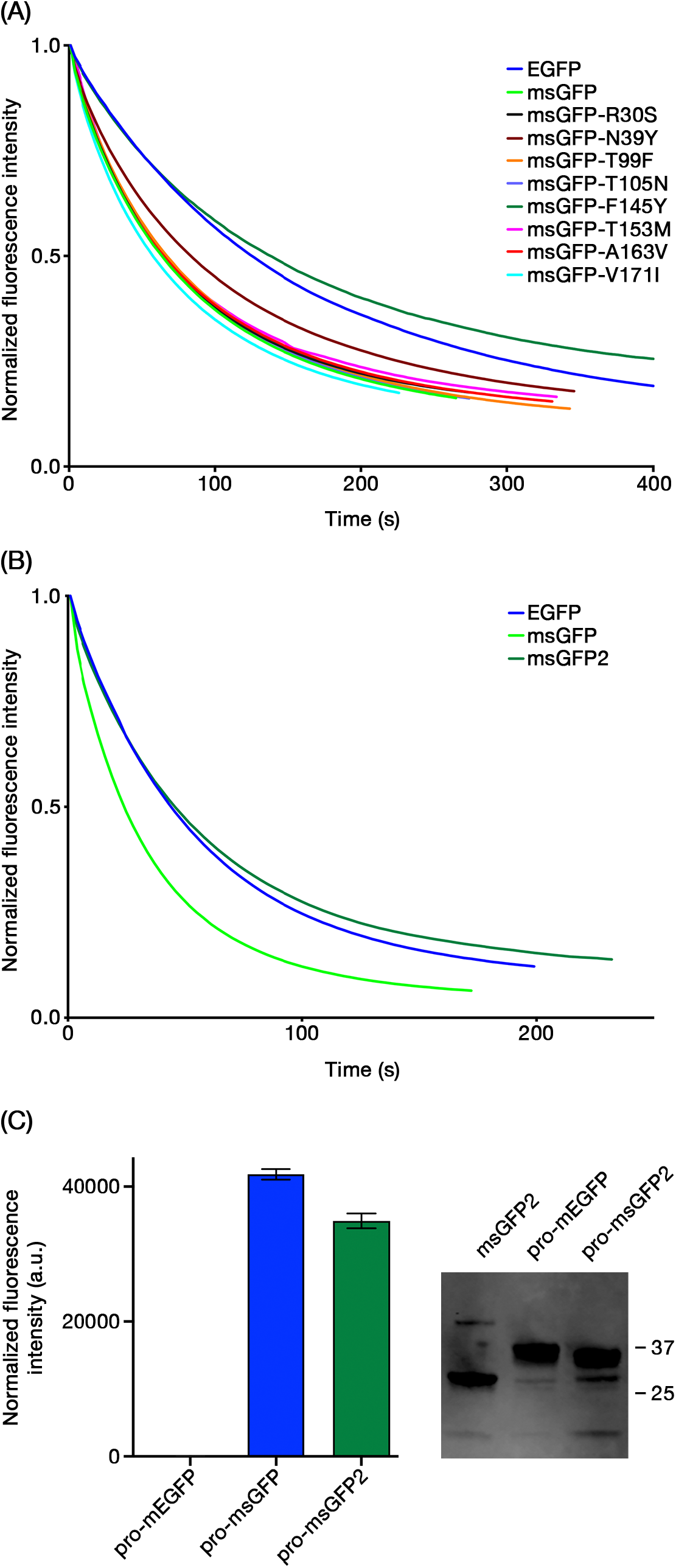
The F145Y reversion in GFP restores photostability without compromising superfolder activity. (A) As described in Methods, EGFP, msGFP, and the indicated point revertants of msGFP were purified and subjected to photobleaching with blue light illumination on a microscope stage. Each signal was normalized to a starting value of 1.0. (B) Photobleaching of purified EGFP, msGFP, and msGFP2 was analyzed as in (A), except that the blue light illumination was stronger, and the curves were obtained by averaging the results from three experiments. (C) As described in Methods, the indicated proinsulin fusion constructs were expressed in *E. coli*, and the green fluorescence signals were measured. The averaged signals from samples analyzed in triplicate are plotted in arbitrary units with SEM. For comparison, constructs expressing non-fused mEGFP or msGFP2 yielded signals of approximately 13,000 or 57,000 arbitrary units, respectively. To the right of the plot is an immunoblot with an anti-GFP antibody showing the levels of proinsulin-mEGFP and proinsulin-msGFP2 in extracts from the bacterial cells, with unfused msGFP2 as a control. The numbers indicate the sizes in kDa of molecular weight markers.

We reasoned that one of the eight superfolder mutations in msGFP might be responsible for the reduced photostability. Those mutations were reverted individually, and the resulting purified FPs were examined. Strikingly, the F145Y reversion restored photostability to a level at least as high as that of EGFP (Figure 2A, dark green curve). None of the other reversions had a comparable effect. The F145Y reversion was therefore incorporated into msGFP2, which was much more photostable than msGFP and at least as photostable as EGFP (Figure 2B). msGFP2 was similar to EGFP not only with regard to photostability, but also with regard to fluorescence spectra and brightness (Table 1).

**TABLE 1.** Photphysical properties msGFP2 compared to EGFP.

| **FP** | **Absorbance Maximum** | **Emission Maximum** | **Quantum Yield** | **Extinction Coefficient** |
| --- | --- | --- | --- | --- |
| msGFP2 | 489 nm | 509 nm | 0.59 | 52,200 |
| EGFP | 488 nm | 510 nm | 0.60 | 52,900 |

To confirm that the F145Y reversion did not abolish superfolder activity, we generated fusions of different GFP variants to mammalian proinsulin, which normally has disulfide bonds and is expected to misfold in the bacterial cytoplasm and form inclusion bodies (20). Expression of a proinsulin-mEGFP fusion in bacteria yielded no detectable fluorescence, presumably because mEGFP cannot mature to the fluorescent state in inclusion bodies (5). By contrast, expression of proinsulin-msGFP yielded strong fluorescence (Figure 2C). With proinsulin-msGFP2, the signal was slightly lower but still strong (Figure 2C). As judged by immunoblot analysis of the bacterial cell extracts, the proinsulin-mEGFP and proinsulin-msGFP2 constructs were expressed at similar levels, indicating that the lack of fluorescence with proinsulin-mEGFP reflected misfolding (Figure 2C). These results show that msGFP2 retains superfolder activity.

In sum, compared to EGFP, the superfolder variant msGFP2 has similar photophysical properties but contains the following changes:

- The monomerizing A206K mutation
- The seven superfolding mutations S30R, Y39N, F99T, N105T, M153T, V163A, and I171V (but not Y145F)
- An N-terminal MDSTES peptide in place of MVSKGEE
- A C-terminal GGSGGS peptide in place of LGMDELYK

### 2.5 Functional tests of msGFP2 in yeast

In the oxidizing environments of the ER lumen and the bacterial periplasm, superfolder GFP variants perform better than traditional GFP variants, which undergo aberrant disulfide bonding (6, 21). We therefore tested whether msGFP2 was less disruptive than mEGFP as a tag for *Saccharomyces cerevisiae* Kar2, the yeast homolog of the ER luminal chaperone BiP (22). Chromosomal gene replacement was used to create Kar2-mEGFP and Kar2-msGFP2 fusions. At 23°C or 30°C, Kar2-msGFP2 exhibited a typical yeast ER pattern consisting of a prominent nuclear envelope ring plus weaker cortical labeling (Figure 3A) (23). By contrast, Kar2-mEGFP produced fluorescent aggregates at both 23°C and 30°C (Figure 3A). When growth was examined at 30°C, the strain expressing Kar2-msGFP2 grew similarly to a strain expressing wild-type Kar2, but the strain expressing Kar2-mEGFP grew more slowly (Figure 3B). These results fit with a previous study that employed a superfolder GFP to tag Kar2 (24), but msGFP2 exhibits photostability that should make it a preferred option for live cell imaging.

**FIGURE 3.**
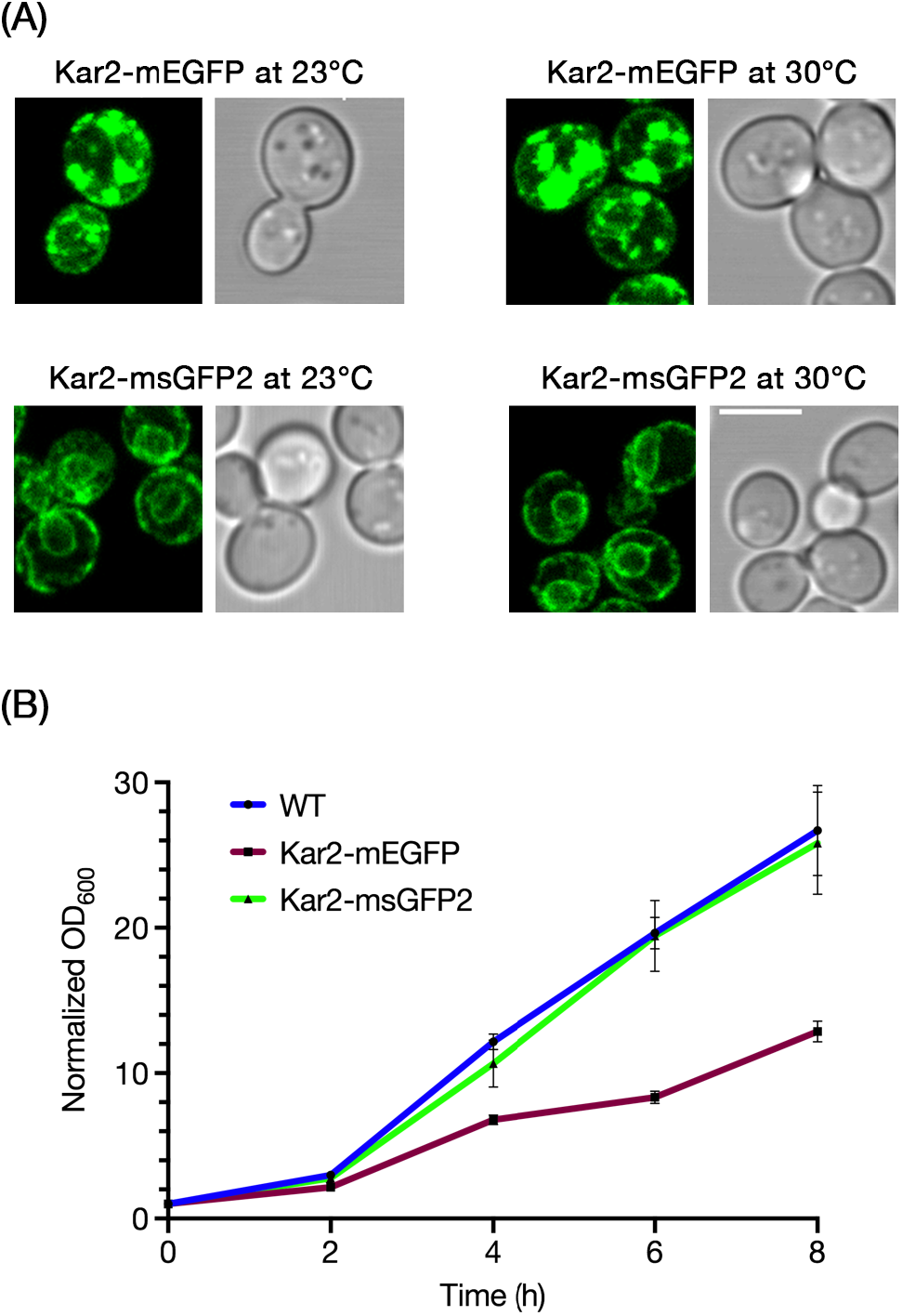
msGFP2 outperforms mEGFP as a fusion tag for the *S. cerevisiae* ER protein Kar2. (A) The endogenous Kar2 protein was tagged with mEGFP or msGFP2 by gene replacement. Cells grown in minimal medium at either 23°C or 30°C were imaged by confocal microscopy. Optical sections corresponding to approximately the central half of each cell were projected. Scale bar, 5 µm. (B) Cells expressing wild-type Kar2, or Kar2-mEGFP, or Kar2-msGFP2 were grown overnight at 30°C in rich medium, then diluted to an OD_600_ of approximately 0.1 in the same medium. The diluted cultures were grown further at 30°C, and OD_600_ values were measured at 2-h intervals. OD_600_ values for each culture were normalized to the starting value. The graph represents averaged values for three independent experiments, with bars indicating SEM.

Are the benefits of msGFP2 also apparent when tagging a cytosolic protein? To address this question, we built on the finding that in the yeast *Pichia pastoris*, tagging the COPII vesicle coat protein Sec13 with GFP results in lethality at 36.5°C when combined with the P1092L point mutation in Sec16 (25). The implication is that the Sec13-GFP fusion protein is partially defective. This effect was seen by plating yeast cells in a dilution series (Figure S2). At 23°C, *sec16-P1092L* mutant cells grew well when Sec13 was tagged by gene replacement with either EGFP, or mEGFP, or msGFP2. By contrast, at 36.5°C, the strain expressing Sec13-msGFP2 grew much better than the strains expressing Sec13-EGFP or Sec13-mEGFP (Figure S2). This effect was quantified by measuring growth in liquid culture at the semipermissive temperature of 34°C. While none of the tagged Sec13 constructs conferred wild-type growth rates, the cells expressing Sec13-msGFP2 grew substantially better than those expressing Sec13-EGFP or Sec13-mEGFP (Figure 4A). We conclude that Sec13 function is sensitive to tagging, and that msGFP2 is less perturbing in this context than standard GFP variants.

**FIGURE 4.**
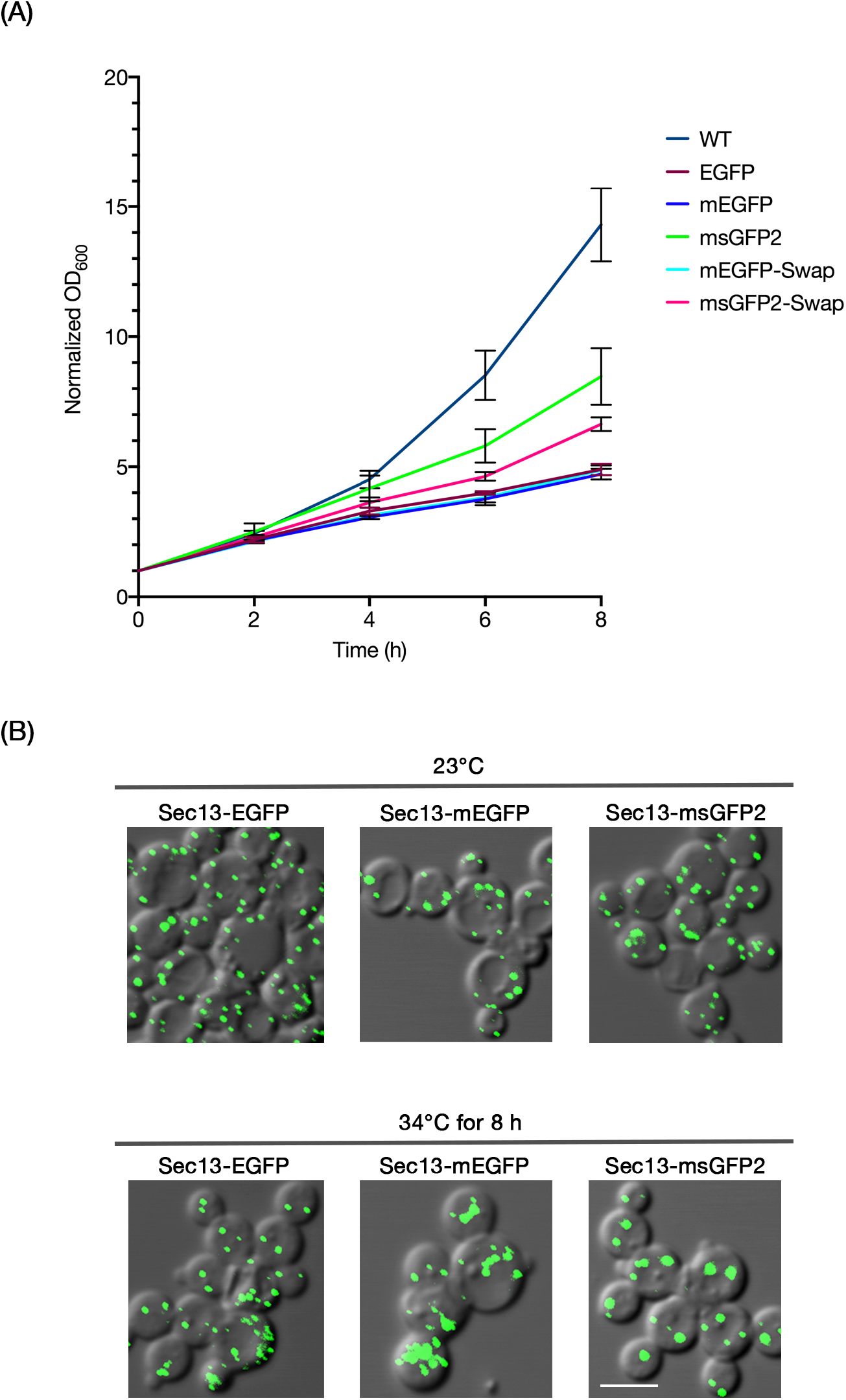
The superfolder property of msGFP2 enhances functionality in a fusion with the *P. pastoris* COPII coat protein Sec13. (A) *P. pastoris* strains with the indicated tags on Sec13 were grown overnight at 23°C in rich medium to early log phase. The cultures were then diluted in rich medium to OD_600_ values of about 0.3, and incubated at 34°C for 8 h. OD_600_ measurements were taken every 2 h for three independent cultures per construct, normalized to the initial OD_600_ values, and averaged. Bars represent SEM. (B) Aliquots of the cells from (A) were imaged by fluorescence microscopy after initial growth at 23°C, and then again after incubation for 8 h at 34°C. Scale bar, 5 µm.

The stronger growth of the Sec13-msGFP2 strain could reflect either the superfolder property of msGFP2, or the modified N- and C-terminal peptides, or both. To test these possibilities, we created a construct in which Sec13 was fused either to mEGFP containing the msGFP2 terminal peptides (Sec13-mEGFP-Swap), or to msGFP2 containing the mEGFP terminal peptides (Sec13-msGFP2-Swap). As judged by plating yeast cells in a dilution series, only the Sec13-msGFP2-Swap construct yielded significant growth at 36.5°C (Figure S2). This growth was reproducibly a bit weaker than that obtained with Sec13-msGFP2, suggesting that the terminal peptides of msGFP2 made a difference. Similar results were obtained by comparing growth of the Sec13-msGFP2 and Sec13-msGFP2-Swap constructs in liquid culture at 34°C (Figure 4A). In both assays, the superfolder property was necessary for improved growth, and the msGFP2 terminal peptides yielded a further improvement. All of the constructs showed qualitatively similar fluorescence at ER exit sites (Figure 4B) (25, 26), suggesting that the relevant effect of the superfolder property may have been the conformational stability of the GFP tag rather than the cellular level of functional Sec13.

## 3 DISCUSSION

Variants of *A. victoria* GFP are typically among the best performing FPs for use as protein tags. Yet some GFP-tagged proteins show mislocalization and/or reduced functionality, suggesting that there is room for further improvement. We describe a variant called msGFP2 that has two enhancements.

The first enhancement in msGFP2 is the F145Y reversion, which restores photostability to the level seen with EGFP. We note that during the engineering of superfolder GFP, Y145F was the only mutation that altered the interior of the protein (5), so it is not surprising that this mutation changed the photophysical properties. Fortunately, Y145F had only a minor effect on folding (5), and we found that this mutation could be reverted with only a slight reduction in the superfolder property. It is likely that other superfolder GFP variants could also benefit from the F145Y reversion.

The second enhancement in msGFP2 is replacement of the N- and C-terminal peptides. These terminal peptides can cause substantial bacterial cytotoxicity when incorporated into other FPs. In particular, the widely used red FP mCherry contains the N- and C-terminal peptides of EGFP (12), and we find that the N-terminal peptide of EGFP can promote cytotoxicity of mCherry during bacterial expression. Better results are seen with an mCherry variant that incorporates an N-terminal peptide related to the one we generated when optimizing tetrameric DsRed (10). This improvement may reflect lower interactivity of the new N-terminal peptide. We also replaced the C-terminal peptide of mCherry, which is MDELYK, with the sequence GGSGGS, based on the expectation that the GFP-derived MDELYK peptide might be prone to undesirable electrostatic and hydrophobic interactions. Similar N- and C-terminal peptide replacements also reduced bacterial cytotoxicity of mScarlet-I (17), a synthetic red FP. We suggest that the common practice of incorporating the EGFP terminal peptides into engineered FPs is unwise, and that the terminal peptides described here are a promising alternative. Because effects of the type seen with red FPs could also occur with GFP in some situations, we incorporated the new N- and C-terminal peptides into msGFP2.

Compared to mEGFP, which has been a favored choice for protein tagging, msGFP2 is expected to be comparable and sometimes superior. The major benefit of msGFP2 is that it offers superfolder behavior without loss of photostability. One example of an environment in which the superfolder property is beneficial is the lumen of the ER. Standard GFP variants can accumulate in the ER in a nonfluorescent state, apparently because cysteines that are normally in the interior of folded GFP can form disulfides that inhibit folding (6, 27). Indeed, we find that when the yeast ER luminal chaperone Kar2 (BiP) is tagged with mEGFP or msGFP2, only the msGFP2 fusion yields an unperturbed ER pattern. Interestingly, Kar2-mEGFP generated aggregates, which must have contained folded mEGFP because they were fluorescent. We speculate that a subset of the mEGFP molecules misfolded to form nonfluorescent aggregates, and that the fluorescent Kar2-mEGFP molecules associated with those aggregates. In any case, Kar2-msGFP was much better behaved than Kar2-mEGFP with regard to both the fluorescence pattern and the ability to support normal cell growth.

The superfolder property can also be beneficial in the cytosol, especially when a tagged protein is in a challenging local environment such as bacterial inclusion bodies (5). Similar challenges may arise whenever a GFP-tagged protein is highly concentrated at a particular cellular location. We explored a test case involving the polymeric coat of COPII vesicles. Our previous work had revealed that when the COPII subunit Sec13 was tagged with EGFP in the yeast *P. pastoris*, the Sec13-EGFP allele rendered the cells thermosensitive for growth when combined with a point mutation in the COPII-associated Sec16 protein (25). The same effect is seen with Sec13-mEGFP. By contrast, Sec13-msGFP2 allows significantly faster growth at elevated temperatures in a strain expressing mutated Sec16. The reason for the better behavior of Sec13-msGFP2 is not obvious, because the fluorescence signals at COPII-containing ER exit sites are similar for Sec13-mEGFP and Sec13-msGFP2. One possibility is that the cells expressing fluorescent Sec13-mEGFP also contain nonfluorescent Sec13-mEGFP molecules, which actively interfere with COPII function. A more intriguing possibility is that conformational stability is important even after a GFP variant has folded to the fluorescent state, with fluorescent Sec13-msGFP2 molecules being more conformationally stable and hence more functional than fluorescent Sec13-mEGFP molecules. Additional work will be needed to clarify how superfolder GFP variants reduce perturbation of tagged proteins.

Diverse applications can be envisioned for derivatives of msGFP2, including variants with altered spectra (1) or with other specialized properties (28). We have obtained good results using a yeast codon-optimized msGFP2 (data not shown). This type of engineering is helping to unlock the full potential of FPs as fusion tags.

## 4 METHODS

### 4.1 Bacterial growth assay

*E. coli* cells of strain DH10B were transformed with pQE-60NA constructs (29) encoding the various FP variants. A transformed culture was diluted 1:30, then spread on an LB + ampicillin plate and grown overnight at 37°C for 19-20 h. The resulting colonies were photographed with a Bio-Rad FX Pro Plus gel imager, using ultraviolet light illumination to detect fluorescent clones with the exposure time adjusted based on FP brightness. These images were processed using ImageJ (30) as follows: images were changed to 8-bit format, inverted to display black colonies on a white background, and subjected to thresholding to remove background spots and nonfluorescent colonies. Colony areas were automatically measured using the “Analyze Particles” tool. For each sample, the areas of at least 200 colonies were averaged.

### 4.2 FP expression and purification

Hexahistidine-tagged FPs were expressed using a pQE-81-based vector (31). For modification of msGFP, the parental construct was subjected to QuikChange mutagenesis to generate single-codon revertants. Constructs were transformed into *E. coli* XL1-Blue cells, and clones were grown overnight in LB + ampicillin. Each culture was then diluted 1:50 in LB + ampicillin, grown to mid-log phase, induced with 1 mM isopropyl β-D-1-thiogalactopyranoside (IPTG), grown overnight, and centrifuged. The pelleted cells were lysed using B-PER II bacterial protein extraction reagent (Thermo Fisher) for 15 min, and centrifuged for 15 min at 27,000xg. The supernatant was adjusted to 300 mM NaCl, vortex mixed, and centrifuged for 1 min at 17,000xg. The soluble extract was incubated with 0.2 mL of Ni^2+^-NTA-agarose slurry (Macherey-Nagel) per 50 mL of starting culture, and mixed end-over-end for 1 h at room temperature. Then the beads were washed three times with 300 mM NaCl, 20 mM imidazole-HCl, pH 7.4, 0.5% Triton X-100, using 2 mL per 50 mL of culture, followed by three more washes with the same buffer lacking Triton X-100. Each FP was eluted by mixing the beads with 0.5 mL of 300 mM imidazole-HCl, pH 7.4 per 50 mL of starting culture for 20 min at room temperature. The final eluate was adjusted to 0.5 mM EDTA and stored in the dark at 4°C.

### 4.3 Photobleaching assay

A purified 6xHis-tagged FP was adjusted to a protein concentration of 60 µM. In parallel, 1% low-melt agarose in 50 mM Na^+^-PIPES, pH 7.0, 100 mM NaCl, 0.02% sodium azide was melted at 50°C, and 500 µL of the melted agarose was supplemented with 15 µL of 2-µm polystyrene beads (Polysciences). Twenty µL each of the FP and agarose/beads solutions were mixed thoroughly, and 2 µL of this mixture was placed on a preheated glass slide and compressed with a preheated coverslip, aiming for a complete and even spread. The polystyrene beads created a layer of uniform thickness. Melted wax was used to seal the edges of the coverslip. After solidification of the agarose, the sample was imaged. To induce photobleaching, the slide was exposed to continuous illumination with blue light using a Zeiss Axioplan 2 microscope with a GFP filter cube. Images of a single focal plane were taken every 3 s until the field of view was almost completely dark. Fluorescence intensity values were obtained from each image series, blank subtracted, normalized, and plotted as a function of time.

### 4.4 Superfolder activity assay

Vectors based on pQE60-NA were used to express human proinsulin with C-terminally fused FPs. These constructs were transformed into *E. coli* XL1-Blue cells, and transformants were grown on LB + ampicillin plates. For each transformant, three clones were picked, inoculated in 5 mL LB + ampicillin medium, and grown at 37°C to an OD_600_ of 0.5. Cultures were diluted 4x in the same medium, and 5 mL were placed in a 50-mL baffled flask and grown to an OD_600_ of 0.5 at 37°C with shaking. Three aliquots from each culture were transferred to a black 96-well plate to measure the background fluorescence from uninduced cells. The cultures were then induced by adding IPTG to 1 mM, and were incubated for 1.5 h. Aliquots from each induced culture were adjusted to an OD_600_ of 0.5 and transferred to a black 96-well plate to measure the cellular fluorescence. After subtracting the background values, the fluorescence values were averaged.

### 4.5 Spectral measurements

Purified 6xHis-tagged FPs were serially diluted, and the absorbance and emission spectra of each dilution were measured using an Agilent 8453 UV-Visible spectrophotometer and a Horiba Fluorolog-3 fluorimeter, respectively. The emission spectrum values were integrated and plotted as a function of the peak absorbance values, and the slopes of the lines were used to calculate the quantum yields relative to EGFP, which was assumed to have a quantum yield of 0.60 (1).

### 4.6 Yeast methods

The parental yeast strains were *S. cerevisiae* JK9-3da (32) and *P. pastoris* PPY12 (33). Yeast proteins were C-terminally tagged with FPs using the pop-in/pop-out method for *S. cerevisiae* (34) or integrative gene replacement for *P. pastoris* (26). Cells were grown in baffled flasks in rich YPD medium or nonfluorescent minimal NSD medium (35).

For fluorescence imaging, strains were grown in liquid NSD to mid-log phase at room temperature. Cells were either attached to the bottom of a confocal dish (MatTek Corporation) coated with concavalin A (36), or compressed beneath a coverslip and imaged on a glass slide. Images were captured with a Leica SP5 confocal microscope, and were processed using ImageJ and Adobe Photoshop.

### 4.7 Software analysis and DNA constructs

Statistical analysis was performed with GraphPad Prism (Insightful Science), and DNA constructs were simulated and recorded using SnapGene (Insightful Science). DNA constructs were propagated in *E. coli* strain XL1-Blue, which does not exhibit cytotoxicity. Relevant constructs will be archived with Addgene.

---

**FIGURE S1.**
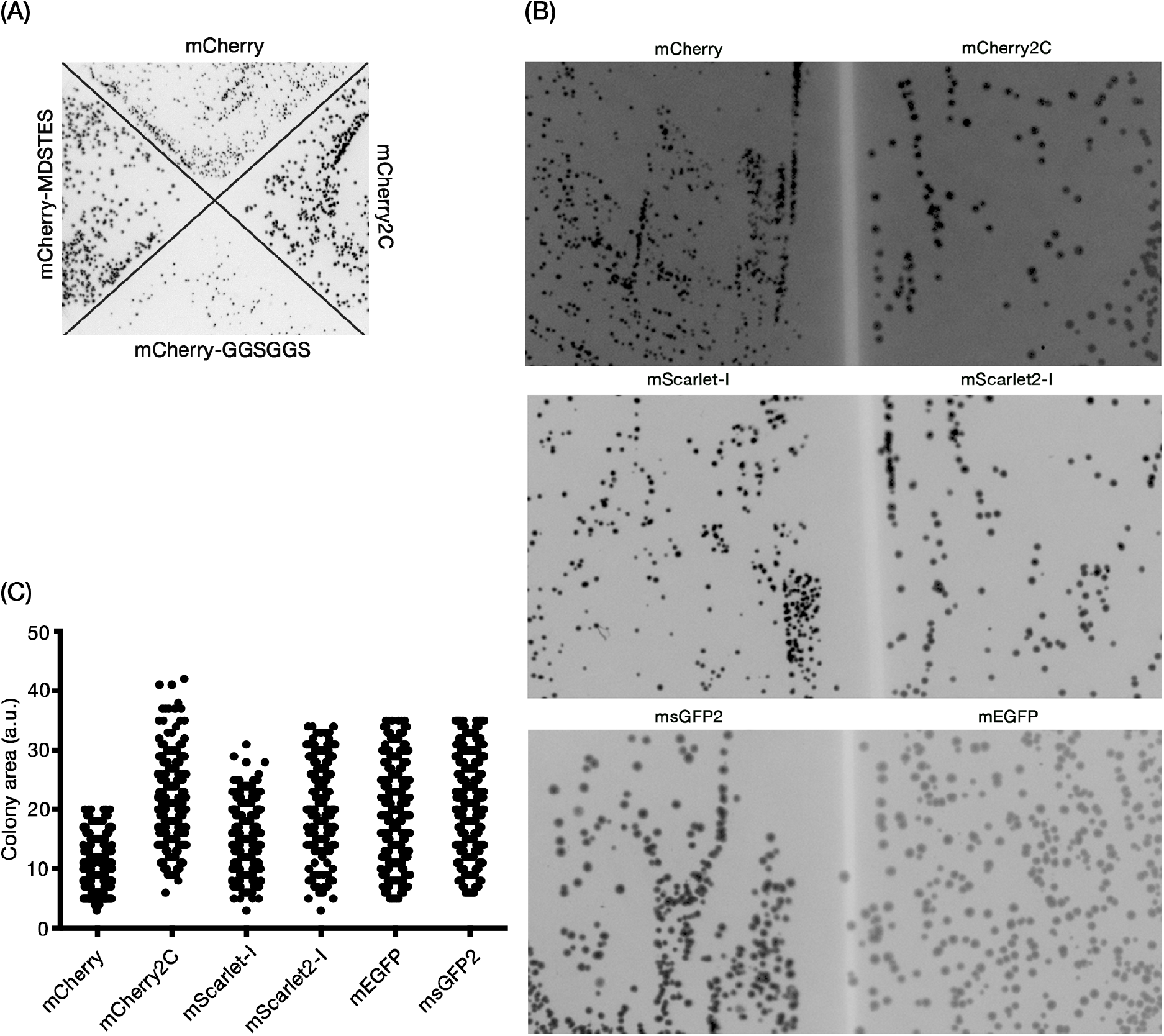
Terminal peptides influence bacterial cytotoxicity for two red fluorescent proteins. (A) *E. coli* DH10B cells were transformed with constructs encoding mCherry, or mCherry2C, or mCherry with its N-terminal peptide replaced with MDSTES, or mCherry with its C-terminal peptide replaced with GGSGGS. Transformants were plated on LB + ampicillin, and the plate was photographed after overnight growth. (B) The experiment was performed as in (A), except that three separate plates were used to compare transformants expressing the indicated pairs of FPs. (C) A scatter plot shows the distributions of colony sizes for the plates in (B).

**FIGURE S2.**
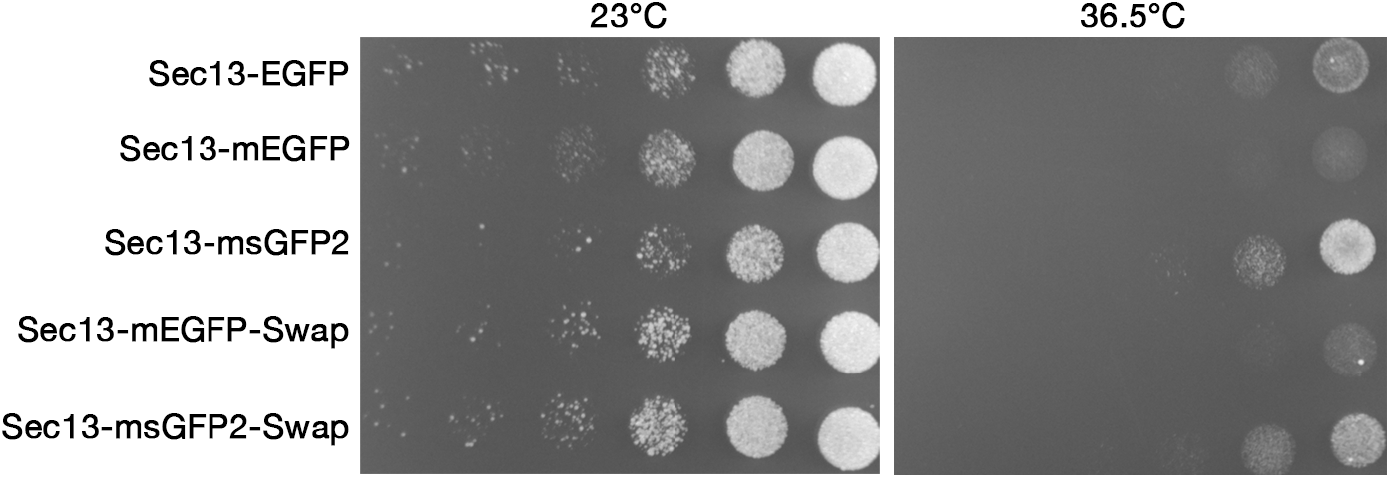
msGFP2 outperforms other GFP variants as a tag for *P. pastoris* Sec13. *P. pastoris* strains with the indicated tags on Sec13 were grown in rich medium to mid-log phase, then diluted to an OD_600_ of 0.25. Serial 10x dilutions were spotted onto two identical minimal medium plates with selection for the integrated tag. The plates were then incubated for 3 days at either 23°C or 36.5°C.

